# Ser149 is another potential 14-3-3 ε binding site of Cdc25B in the G2/M transition of mouse fertilized eggs

**DOI:** 10.1101/2023.08.15.553381

**Authors:** Wen-Ning He, Hai-Yao Pang, Yan-Jun Hou, Shao-Qing Feng, Hui-Ling Zhang, Wen-Xiu Guo, Ru Liu, Jun Meng

## Abstract

Cell cycle division 25B (CDC25B) belongs to the family of cell cycle regulatory proteins. It drives G2/M transition by activating cyclin-dependent protein kinases (CDK1), also known as CDC2, whose activity is directly related to its subcellular localization and phosphorylation state.14-3-3 (YHWA) regulates cell division cycle by binding to Cdc25B as a chaperone protein in mammals. Previously, we found that Cdc25B-Ser149 plays an important role in G2/M transition of mouse fertilized eggs, but the molecular mechanism of this transition remains unclear. In this study, we assessed the role of 14-3-3ε (YHWAE) interaction with phosphorylated Cdc25B-Ser149 in G2/M transition of mouse fertilized eggs. Co-expression of Cdc25B-Ser149A and 14-3-3ε could effectively activate maturation promoting factor (MPF) through direct dephosphorylation of Cdc2-Tyr15, and induce G2 fertilized eggs to enter mitosis rapidly. However, co-expression of the phosphomimic Cdc25B-Ser149D or Cdc25B-WT and 14-3-3ε showed no significant difference in comparison with control groups. 14-3-3ε binds to Cdc25B-WT, which is abolished when Ser149 is mutated to Ala. In addition, we found that 14-3-3ε and Cdc25B were co-localized in the cytoplasm at the G1, S and early G2 phases. Cdc25B was translocated from the cytoplasm to the nucleus at the late G2 phase. However, when Ser149 is mutated to Ala, the cytoplasmic localization of Cdc25B is completely abolished. Our findings suggest that Cdc25B-Ser149 is another specific binding site for 14-3-3ε in G2/M transition of one-cell fertilized mouse eggs, which plays essential roles in the regulation of early development of fertilized mouse eggs.

## Introduction

Cyclin-dependent kinases (CDKs), a kind of serine/ threonine kinase proteins, are central regulators of eukaryotic cell division cycle and G2/M transition and can bind to cyclins to drive cell cycle progression [1–4]. Cdc25B gene is located in autosomal 20p13 and encodes a protein composed of 580 amino acid residues. It is one of the isomers of Cdc25, a member of the protein tyrosine phosphatase family that regulates the cell cycle, and plays a role at specific stages of the cell cycle[5]. Cdc25B activates the Cdc2-CyclinB complex (MPF) by dephosphorizing the Thr14 and Tyr15 of Cdc2 (CDK1), promotes the transition of G2/M phase, and positively regulates the cell cycle process[6]. PKA is a cAMP-dependent ser/thr kinase that phosphorylates Cdc25B-Ser149 and Ser321 in the G1, S and early G2 to regulate cell cycle division, growth, differentiation and proliferation[7, 8]. Cdc25B is dephosphorylated in late G2 and directly dephosphorylates Cdc-Tyr15 to restore mitosis in fertilized eggs[9]. The above studies support that Cdc25B plays an essential regulatory role in G2/M transitions in mitosis[10, 11]. The 14-3-3 family consists of phosphoserine/threonine-binding proteins that act as molecular chaperones to regulate the expression of downstream target proteins by binding to ligands containing phosphoserine/threonine sites and its molecular weight is 25-33KDa [12–15]. This regulatory function is a wide range of cellular phenomena, including metabolism, gene regulation, cell cycle, differentiation, migration and apoptosis [16]. At least seven known isoforms of 14-3-3 protein are expressed in mammals: β, ε, η, γ, ζ and σ and τ. Besides, there are two isoforms in yeast, drosophila and nemathelminth [17–19]. The support of some research results laid the foundations for our further research. For instance, 14-3-3ε binds to Cdc25A to activate Akt/BAD/Survivin anti-apoptotic signaling in skin cancer cells [20]. Studies have shown that in Xenopus oocytes, 14-3-3 protein binds to phosphorylated Cdc25C-Ser287 by PKA to regulates oocyte maturation[21]. Uchida S also demonstrated that 14-3-3β, ε binds to phosphorylated Cdc25B-Ser309 in human cells which drives Cdc25B to localize to the cytoplasm[22]. Cui and his colleagues prove that when the endogenous 14-3-3ε is knocked down by siRNA, the cell cleavage rate of fertilized mouse eggs will decreased and cause high abnormal cleavage this result is the same as cardiomyocyte [7, 23]. In our former study, it has been demonstrated that Cdc25B-Ser321 (equivalent to Ser323 in HeLa cells) is a primary 14-3-3ε binding site and this binding blocks the catalytic site of Cdc25B to inhibit the function of the Cdc25B in fertilized mouse egg, thereby directly inhibiting cell division[7]. 14-3-3ε and Cdc25B-Ser321 play a critical regulatory role in the development of mouse embryos by modification of phosphorylation and dephosphorylation, whereas the Ser229 of Cdc25B has no effects on the development of fertilized mouse eggs [24, 25]. Mouse fertilized egg is a cell cycle model similar to that of human species in vertebrates.Little information was found about the molecular mechanism of 14-3-3ε and Cdc25B-Ser149 in the early development of mouse embryos, in particular about that how to regulate the G2/M transition. Besides, there is no evidence to prove whether Ser149 is potential binding site of 14-3-3ε to regulate mitosis in fertilized mouse eggs.

In this research, we investigated seven isoforms of 14-3-3 protein and find that 14-3-3ε is a crucial role in regulating cell division of fertilized mouse eggs. And our data suggest that Cdc25B-Ser149 (corresponding to Ser151 in the human protein) regulates the development of fertilized mouse eggs and Ser149 in mammalian cells, may be potential binding sites of 14-3-3ε involved in mitosis of fertilized mouse eggs. It plays an important role in G2/M transition of fertilized mouse eggs and the subcellular localization of Cdc25B. Our results further prove that phosphorylation of Cdc25B-Ser149 which can be phosphorylated by PKA is a prerequisite for binding to 14-3-3ε which could regulate both function and localization of Cdc25B and indirectly regulates the early development of mouse embryos.

## Materials and Methods

### Animal

Kunming strain mice which are genealogy-specific and pathogen-free (females are 4 weeks age and 20g weight and males are 8 weeks age, 30g weight) were obtained from the Department of Laboratory Animals, Inner Mongolia University (IMU). All experiments were performed at Inner Mongolia Medical University (IMMU) in accordance with the National institutes of Health guidelines for the Care and Use of Laboratory Animals. The protocol for animal handling and the treatment procedures were reviewed and approved by the IMMU Animal Care and Use Committee. Reagents, unless otherwise specified, were from Sigma.

### Collection and culture of mouse embryos

One-cell stage mouse embryos were collected and cultured according to the method described in our previous report [26] based on Hogan and Constantini’s method [20]. The embryos were incubated in the M16 medium containing 200μmol/L PKA activator, dibutyryl cAMP (dbcAMP) covered with mineral oil at 37℃, 5% CO_2_ in the air after injection with various mRNA or plasmids.

### RNA extraction, cDNA synthesis and real-time PCR

Real-time PCR mRNAs were extracted from G1, S, G2 and M phases of fertilized mouse eggs using the EllustraTM QuickPrep MicromRNA Purification Kit (GE healthcare UK Limited, UK). The cDNA is synthesized using Prime Script RT reagent Kit with gDNA Eraser (TakaRa). The RT reaction was carried out for one cycle at 42℃ for 30 min; 99℃ for 5 min; 4℃ for 5 min. The real-time PCR reaction was performed using One Step SYBR Prime Script RT-PCR Kit (TakaRa). The primers for 14-3-3β, 14-3-3γ, 14-3-3ε, 14-3-3ζ, 14-3-3η, 14-3-3σ, 14-3-3τ and β-actin were designed and synthesized by Takara (Table 1). To run the real-time PCR reaction, the following protocol was followed: 95℃ for 30 sec, 95℃ for 3 sec, 60℃ for 30 sec (40 cycles), 95℃ 15 sec, 56℃ for 30 sec, 95℃ for 15 sec (one cycle).

**Table 1.**
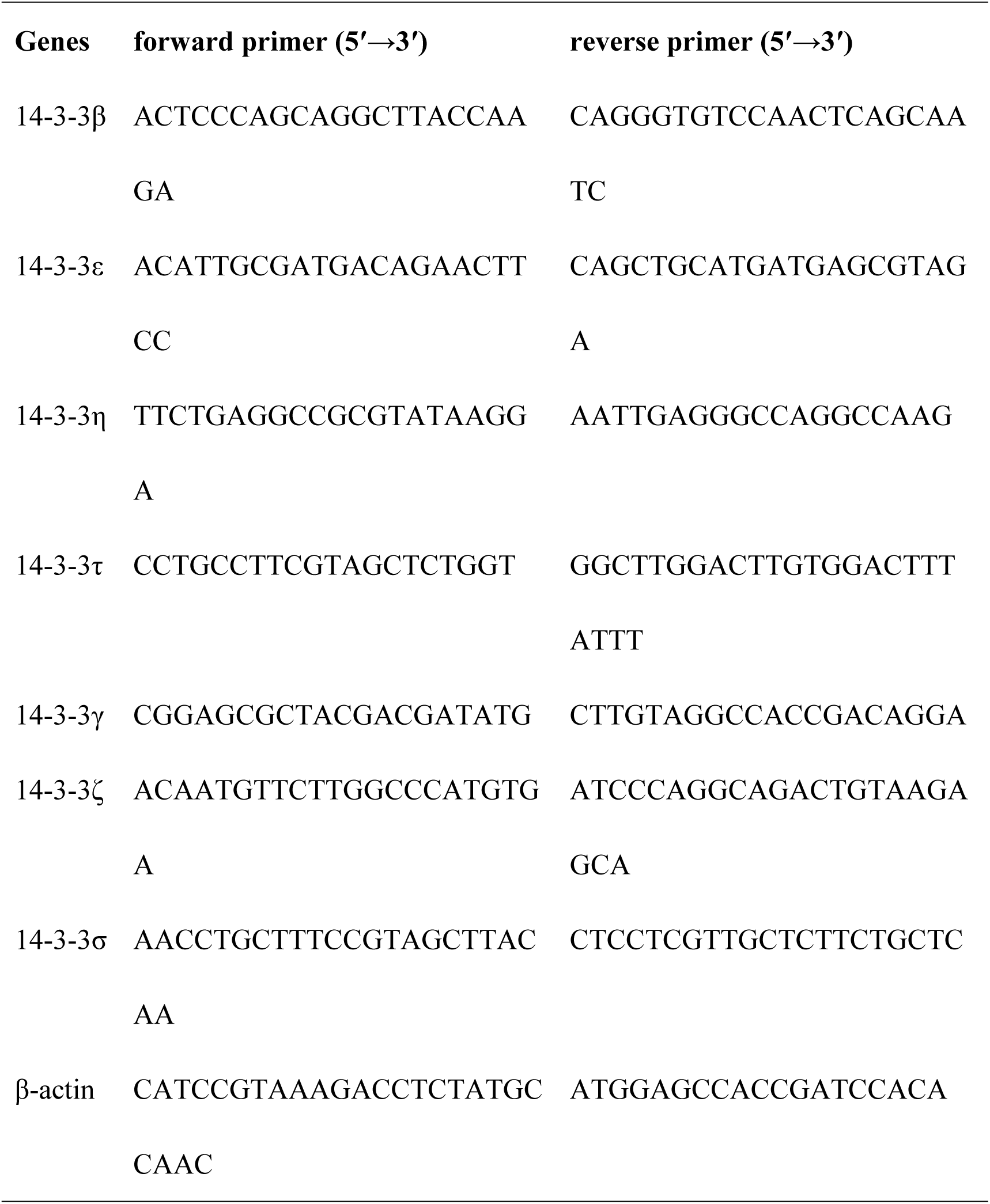
Primers for real-time PCR.

### Immunofluorescence

Embryos in G1, S, G2 and M phases were washed in phosphate-buffered saline (PBS) containing 0.1% bovine serum albumin (BSA) and fixed in 4% paraformaldehyde in PBS (pH=7.4) at room temperature for one hour. After being permeablilzed with 0.1% TritonX-100 in PBS at room temperature for 30 min, embryos were blocked in 5% BSA in PBS for 1 h and incubated overnight at 4℃ with polyclonal goat anti-Cdc25B antibody diluted (1:100, Santa Cruz Biotechnology) and polyclonal rabbit anti-14-3-3ε antibody diluted (1:800, Abcam) at 4℃. Following three washes in a solution consisting of PBS with 0.1% BSA, the embryos were incubated with FITC-conjugated goat anti-rabbit secondary antibody (dilution1:50 in PBS) and TRITC-conjugated rabbit anti-goat secondary antibody (dilution 1:50 in PBS) at 37℃ for 1 h. Then, the DNA was stained with 25μg/ml Hoechst33258 for 10 min at room temperature. The signals of subcellular localization of endogenous Cdc25B and 14-3-3ε and DNA staining were detected by a Laser Confocal Scanning Microscope at 488 nm, 530 nm and 260 nm, respectively.

### In Vitro Transcription

As described in our previous report [18], the pcDNA3.1-ZEO-HA-14-3-3ε and all of the pcDNA3.1-Myc-Cdc25B constructs were linearized with XbaI and transcribed in vitro into 5**^’^**-capped mRNA for microinjection using a mMESSAGE mMACHINE kit (Ambion).

### Microinjection and Morphology Analysis

Various Cdc25B or 14-3-3ε plasmids and 14-3-3ε or Cdc25B mRNAs were microinjected into the nucleus or cytoplasm of one-cell embryos at G1 or S phases according to our previous report [26]. Typical injection volume was 5% (10 pl, cytoplasm) and 1% (2 pl, nucleus) of total cell volume per egg. Messenger RNAs were diluted to various concentrations in TE buffer (5 mM Tris-HCL and 0.5 mM EDTA, pH=7.4), respectively, without nuclease contaminant. Eggs in control groups were either not microinjected or microinjected with TE buffer.

Mitotic stages (G1, S, G2 and M phases) were defined as previously described [20]. The cleavage rate, i.e. the number of two-cell embryos resulting from one-cell embryo division, was counted from three independent experiments under a phase-contrast microscope 31 or 34 hours after hCG injection in the presence of 2 mmol/l dbcAMP in control and microinjection groups. Morphological analysis was performed by inverted microscope.

To observe the subcellular co-localization of various Cdc25B and 14-3-3ε, plasmid DNA of pEGFP-Cdc25B-WT, pEGFP-Cdc25B-Ser149A were co-injected with pmax-RFP-HA-14-3-3ε into the nucleus of mouse fertilized eggs at G1 phase at the concentration of 1μg/μl. After microinjection the embryos were transferred from M2 medium to M16 medium containing 2 mmol/l dbcAMP until a distinct fluorescent signal was detected.

### MPF activity assay

Histone H1 kinase assay was employed to measure the activity of MPF kinase [21]. Five fertilized eggs cultured in M16 medium were collected, washed in collection buffer (PBS containing 1 mg/ml polyvinyl alcohol, 5 mM EDTA, 10 mM Na_3_VO_4_, and 10 mM NaF), and then transferred to an Eppendorf tube containing 5 μl of the collection buffer. The Eppendorf tube was immediately stored at -70℃ until the kinase assay was performed. An identical procedure as in previously published reports [9, 26] was followed for the kinase activity assay.

### Western blotting

Oocytes were lysed at the indicated time points, subjected to sodium dodecyl sulfate-polyacrylamide gel electrophoresis (SDS-PAGE) (12%), and subsequently transferred onto nitrocellulose membranes. Blots were blocked for 1 h at room temperature in 5% milk in Tris buffered saline (TBS) containing 0.05% Tween20 (TBST). The blots were probed overnight at 4℃ in 1% milk in TBST with the following antibodies: rabbit anti-14-3-3ε (1:1000) (Abcam); rabbit anti-Myc (1:1000) (Clontech); rabbit anti-HA (1:800) (Sigma); Tyr(P)15 of Cdc2 (1:500; Santa Cruz Biotechnology) and rabbit anti-β-actin (1:400) (Santa Cruz Biotechnology). For detection, the HRP-conjugated anti-rabbit or anti-goat secondary antibody at a dilution of 1:5000 was employed (Beijing Zhongshan Biotechnology, China). Proteins were visualized using the enhanced chemiluminescence (ECL) detection system (Pierce Biotechnology). The proteins expression of 14-3-3ε and β-actin were detected by Western blotting. Densitometry of bands was performed with Quantity One Software, densitometry of 14-3-3ε bands/densitometry of β-actin bands was used as quantitation of endogenous 14-3-3ε expression. Bars represent means ±S. D of three independent experiments.

### Cell culture and transfection for Immunoprecipitation

HEK293 cells (ATCC number CRL-1573) were cultured in Dulbecco’s Modified Eagle’s medium (DEME) (Sigma) supplemented with 10% fetal bovine serum (FBS) (Invitrogen)),100 U/ml penicillin and 10 µg/ml streptomycin. Transient transfections were performed with FuGENE6 (Roche Molecular Biochemicals, Germany) according to the manufacturer’s instructions. For immunoprecipitation, 2 million cells were typically seeded at each well. After culturing overnight, cells were co-transfected with 5µg of pcDNA3.1-flag-Cdc25B -WT, Ser149A, Ser149D or pcDNA3.1-flag Vector and 5 mg of pmax-RFP-myc-14-3-3ε DNA. Transfected cells were processed for immunoblotting or immunoprecipitation after 48h.

Preparation of crude cell extracts, immunoprecipitation and immunoblotting Transfected cells were lysed in immunoprecipitation (IP) buffer (50 mM Tris-HCl, 0.25%w/v Deoxycholate, 1% NP40, 150 mM NaCl, 1 mM EDTA, pH 7.4) supplemented with a protease inhibitor mix and a phosphatase inhibitor mix. The protease inhibitor mix contained 5 µg/ml leupeptin, pepstatin A, aprotinin, and 1mM PMSF. The phosphatase inhibitor mix consisted of a 1:100 dilution of Phosphatase inhibitor cocktail II (Sigma, USA), 25 mM NaF, 0.1 mM sodium orthovanadate and 25 mM β-glycerophosphate. Cell lysates were incubated with mouse monoclonal anti-MYC-antibody (Covance) and protein G-agarose beads (GE Healthcare). The precipitates were then washed with a lysis buffer. Cell lysates and immunoprecipitates were analyzed on Western blots using mouse monoclonal anti-MYC antibody (1:800; Covance) (for exogenous 14-3-3ε) or anti-FLAG antibody (1:800; Sigma) (for exogenous Cdc25B) and anti-GAPDH antibody (1:4000).

### Statistical analysis

Differences in the rate of cleavage of one-cell embryo microinjected with different mRNAs were evaluated using the Chi-square test. Differences in the MPF activity assay were evaluated by one-way analysis of variance followed by a Least Significant Difference (LSD) test. A P value less than 0.05 indicates a significant difference.

## Results

### Identification of 14-3-3 isoforms expressed in fertilized mouse eggs

To explore the expression profile of 14-3-3 isoforms in fertilized mouse eggs at G1, S, G2, M phases, 200 eggs from different phases were respectively collected and used to amplify the mRNA of seven 14-3-3 isoforms. The results of RT-PCR indicate that all of the seven isoforms are expressed in four phases of fertilized mouse eggs. However, their relative expression of mRNA is different at different phase (P<0.05). Relative expression of 14-3-3ε and 14-3-3ζ mRNA were higher than that of other five isoforms in four phases (Figure 1A).

**Figure 1.**
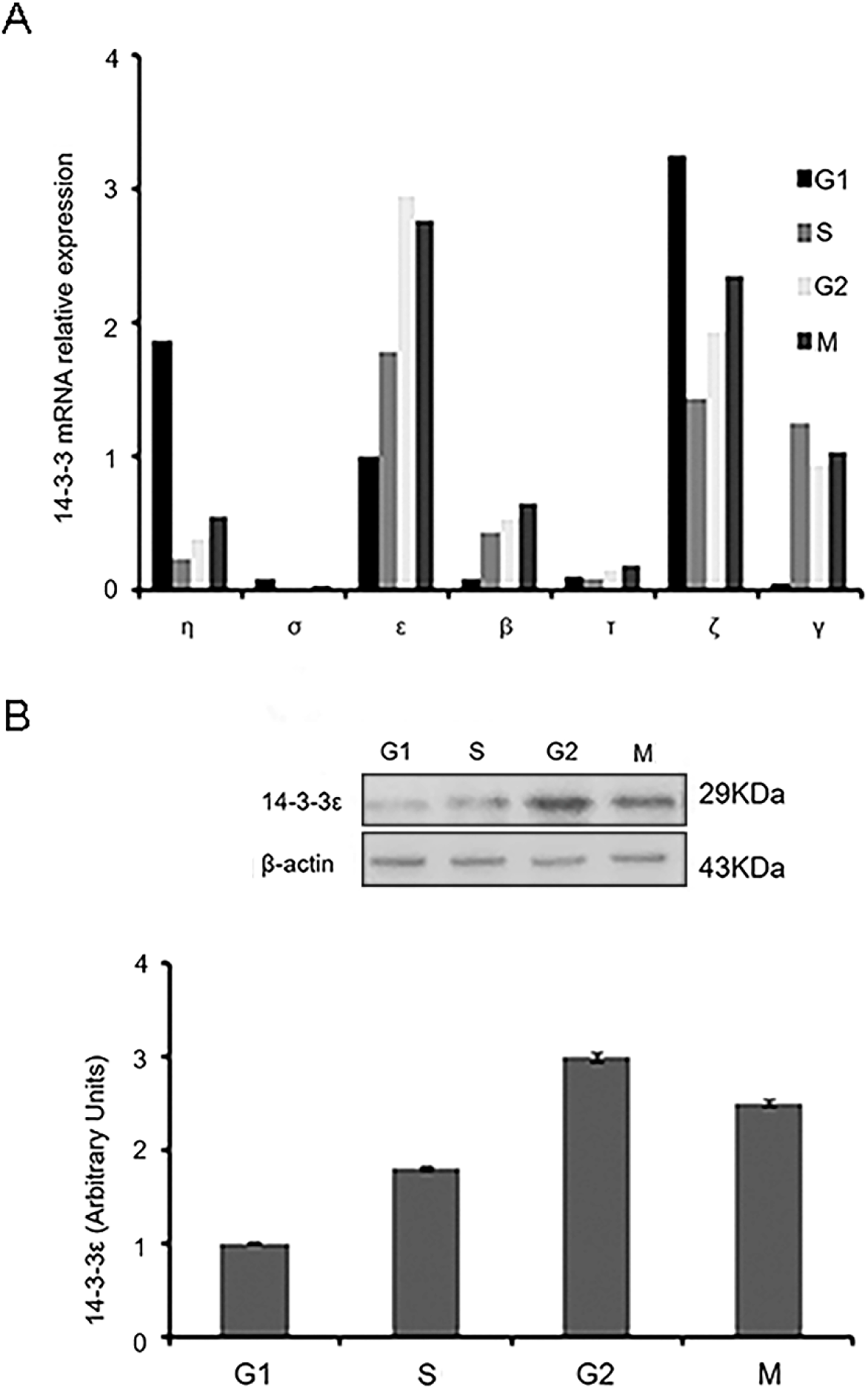
Identification of 14-3-3 isoforms expressed in fertilized mouse eggs. (A) The mRNA relative expression levels of 14-3-3 isoforms at G1, S, G2, M phases of mouse fertilized eggs. The mRNA of 150 fertilized mouse eggs was extracted in G1, S, G2, M phases. RT-PCR products using primers for specific 14-3-3 isoforms are observed in Real time-PCR. 14-3-3ε, 14-3-3β, 14-3-3γ, 14-3-3ζ, 14-3-3η, 14-3-3σ, and 14-3-3τ represent seven 14-3-3 isoforms. (B) Western blot analysis of 14-3-3ε and β-actin protein expression with an anti-14-3-3ε antibody or anti-β-actin antibodies at G1, S, G2, M phases and 300 fertilized eggs are loaded onto each lane(P<0.05). Line G1: fertilized mouse eggs at G1 phase, Line S: fertilized mouse eggs at S phase, Line G2: fertilized mouse eggs at G2 phase, Line M: fertilized mouse eggs at M phase. 14-3-3ε protein band of approximately 29 KDa (upper panel). β-actin protein band of approximately 43 KDa (lower panel). Densitometric quanitification represents the protein expression of 14-3-3ε at G1, S, G2, M phases (column chart) (P<0.05). A *x*^2^ test was used to evaluate the differences of endogenous 14-3-3ε mRNA expression levels or protein expression of 14-3-3ε between multiple experimental groups. Bars represent means ± S.D of three independent experiments.

The expression of 14-3-3ε gradually increased from G1 to G2 and decreased slightly in M phase. We also examined the protein expression of 14-3-3ε by Western blotting and the result demonstrated that the protein expression pattern of 14-3-3ε was consistent with its mRNA expression (Figure 1B). Santanu De and his colleagues [27] have reported that mouse mature metaphase II-arrest eggs express all seven 14-3-3 isoforms and 14-3-3β, 14-3-3ε,14-3-3η and 14-3-3ζ appear in lesser amounts in mature metaphase II-arrest eggs than in immature oocytes which is consistent to our results. Furthermore, it has proved that only 14-3-3ε expressed in GV and GVBD mouse oocytes which is one of the seven 14-3-3 isoforms and the expression of 14-3-3ε remained unchanged during GV and GVBD stages [28].

### Cdc25B-Ser149A, Cdc25B-Ser149D or Cdc25B-WT mRNA interferes with cell division in one-cell mouse embryos

Several reports have demonstrated that dbcAMP has play a critical role in regulating meiotic arrest in mouse fertilized egg and the concentration of 2mmol/l dbcAMP lead to maximal G2 arrest in fertilized mouse eggs[9, 29]. Besides the study also demonstrated that Cdc25B-Ser149A could effectively reverse G2/M arrest can bypass the inhibitory effect of PKA on meiotic resumption induced by dbcAMP.

The Cdc25B-Ser149A mutant had a more potent effect on G2/M transition than Cdc25B-WT, while the Cdc25B-Ser149D mutant displayed similar activities to Cdc25B-WT. In order to verify whether the Cdc25B-Ser149 site can be combined with 14-3-3ε to affect the mitosis progression, one-cell stage fertilized mouse eggs (S phase, 21 h after the hCG injection) firstly were incubated in M16 medium containing 2mmol/l dbcAMP and 1h later microinjected with 14-3-3ε mRNA solely or co-injected with

Cdc25B-WT mRNA, Cdc25B-Ser149A mRNA or Cdc25B-Ser149D mRNA at a concentration of 300μg/ml which were transcripted in vitro. After microinjection, the mouse embryos were transferred back into the M16 Medium containing 2mmol/l dbcAMP, no injection and TE injection groups served as the negative control group . In the group of which were co-injected with 14-3-3ε mRNA and Cdc25B-Ser149A mRNA, the percentage of cleavage of the one-stage mouse embryos is 94.0% which means these fertilized mouse eggs had enter the M phase of mitosis at 34h after hCG injection (12h after the microinjection) (Figure 2). It significantly increased much more than the other group (P<0.05). However, no embryo co-injected with 14-3-3ε mRNA and Cdc25B-WT mRNA or Cdc25B-Ser149D in the presence of 2mmol/l dbcAMP has developed into the two-cell stage until at least 34 h after hCG injection (12h after vary microinjection), which was similar to the control group (no injection and TE injection). In addition, mouse embryos with sole injection with 14-3-3ε mRNA still were arrested at G2 phase at 34h after hCG injection (12h after the microinjection), which implies there is no help of accelerating the process of mitosis with the presence of dbcAMP by overexpression of 14-3-3ε solely (Figure 2). Less than 4% of fertilized mouse eggs were dead after the various injections (p>0.05).

**Figure 2.**
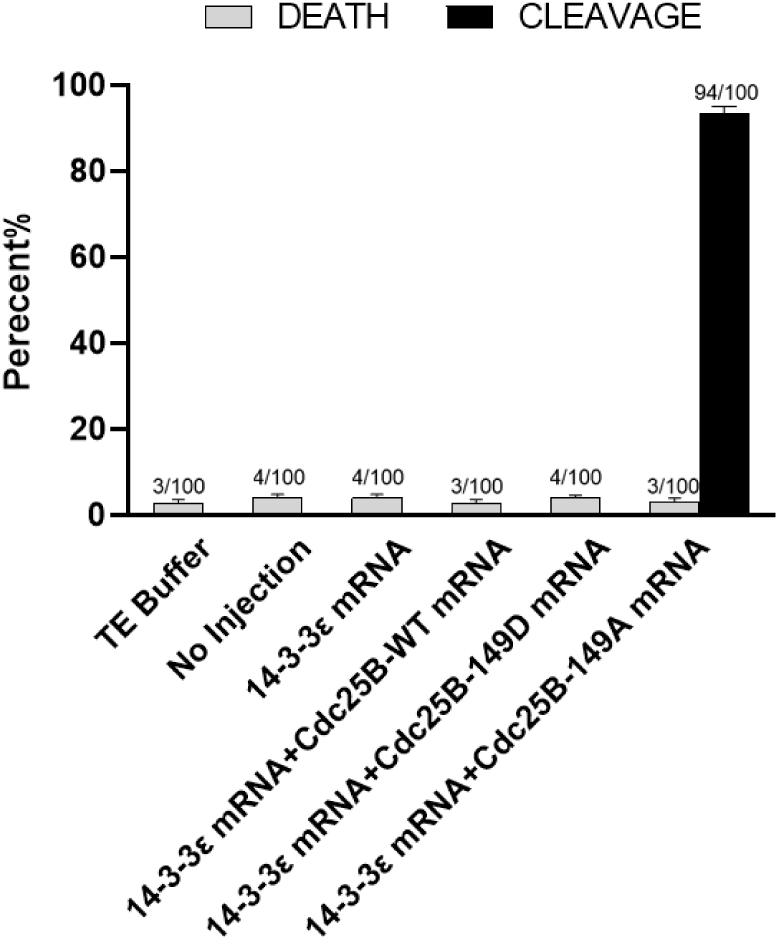
Cell division rate in cultured mouse embryos after Cdc25B-Ser149A or Cdc25B-Ser149D treatment. Effects of microinjection with 14-3-3ε mRNA solely or co-injection with Cdc25B-WT mRNA, Cdc25B-Ser149A mRNA or Cdc25B-Ser149D mRNA and 14-3-3ε mRNA on mitotic resumption of mouse fertilized eggs. Experiments were performed in the continued presence of 2mmol/l dbcAMP. The cleavage rate in cultured mouse embryos of various mRNA microinjection in the presence of 2mmol/l dbcAMP at 34 h after hCG injection. The cleavage rates were calculated with data from three independent experiments. A *x^2^* test was used to evaluate the differences between multiple experimental groups and 300 eggs are calculated for each group. Bars represent means ± S.D of three independent experiments.

### Co-injection of 14-3-3-ε mRNA and CDC25B-Ser149A mRNA overcomes G2 arrest by dbcAMP

To further illustrate our conclusion, we also detected the MPF activity with H1 as the substrate and phosphorylation status of Cdc2-Tyr15 in embryos injected with various mRNAs in the presence of dbcAMP at 29 h after hCG injection every 30 min. In the group of co-injection of 14-3-3ε and Cdc25B-Ser149A mRNA, MPF activity of mouse embryos fluctuated during the maturation process and initially raise at 30h after hCG injection (8h after microinjection) and the peak of MPF activity appear at 9h after microinjection and then began to slowly decline (Figure 3F). This MPF activity differed significantly from the other injected groups (P<0.05). On the contrary, MPF activity consistently maintain a steady low level in the respective group of sole injection of 14-3-3ε mRNA or co-injection of 14-3-3ε and Cdc25b-WT mRNA or 14-3-3ε and Cdc25b-Ser149D mRNA or two controls at 29-32h after hCG injection (7-10h after microinjection) (Figure 3A-E). At the same time, we also measured the phosphorylation status of Cdc2-Tyr15 in the control and microinjection groups by Western blot. In control groups, there strong inhibitory phosphorylation of Cdc2-Tyr15 was observed at 29-30h after hCG injection. It is similar to the group of sole injection of 14-3-3ε mRNA or co-injection of 14-3-3ε and Cdc25b-WT mRNA or 14-3-3ε and Cdc25b-Ser149D mRNA groups. In contrast, in the group co-injected with Cdc25B-Ser149A and 14-3-3ε, a strong signal of Cdc2-Tyr15 phosphorylation was observed only 29 h after hCG injection, and no phosphorylation signal was detected again at 30 h and thereafter (Figure 3G). These change tendency of Cdc2-Tyr15 phosphorylation in each group were consistent with the MPF activity. Moreover, various Cdc25B-MYC fusion proteins and HA-14-3-3ε fusion protein at the protein levels were probed by Western blot in various mRNA-injected fertilized eggs 4h after microinjection (Figure 3H, I). These data indicated that all injected mRNAs were efficiently translated and exogenously expressed protein levels are higher than endogenously expressed protein levels (P<0.05). Furthermore, to determine whether 14-3-3ε can bind Cdc25B at Ser149, HEK293 cells were transfected with myc-14-3-3ε together with empty vector, pcDNA3.1-flag-Cdc25B-WT, flag-Cdc25B-Ser149A or flag-Cdc25B-Ser149D. Recombinant protein expression was confirmed in lysates from transfected cells. 14-3-3ε was immunoprecipitated with anti-myc beads followed by Western blotting to detect Cdc25B binding with 14-3-3ε. However, Cdc25B-WT and Cdc25B-Ser149D were found in a complex with 14-3-3ε, whereas Cdc25B-Ser149A was not detected (Figure 4). This indicates that 14-3-3ε physically interacts with phosphorylated Cdc25B at Ser149, but not with unphosphorylated Cdc25B.

**Figure 3.**
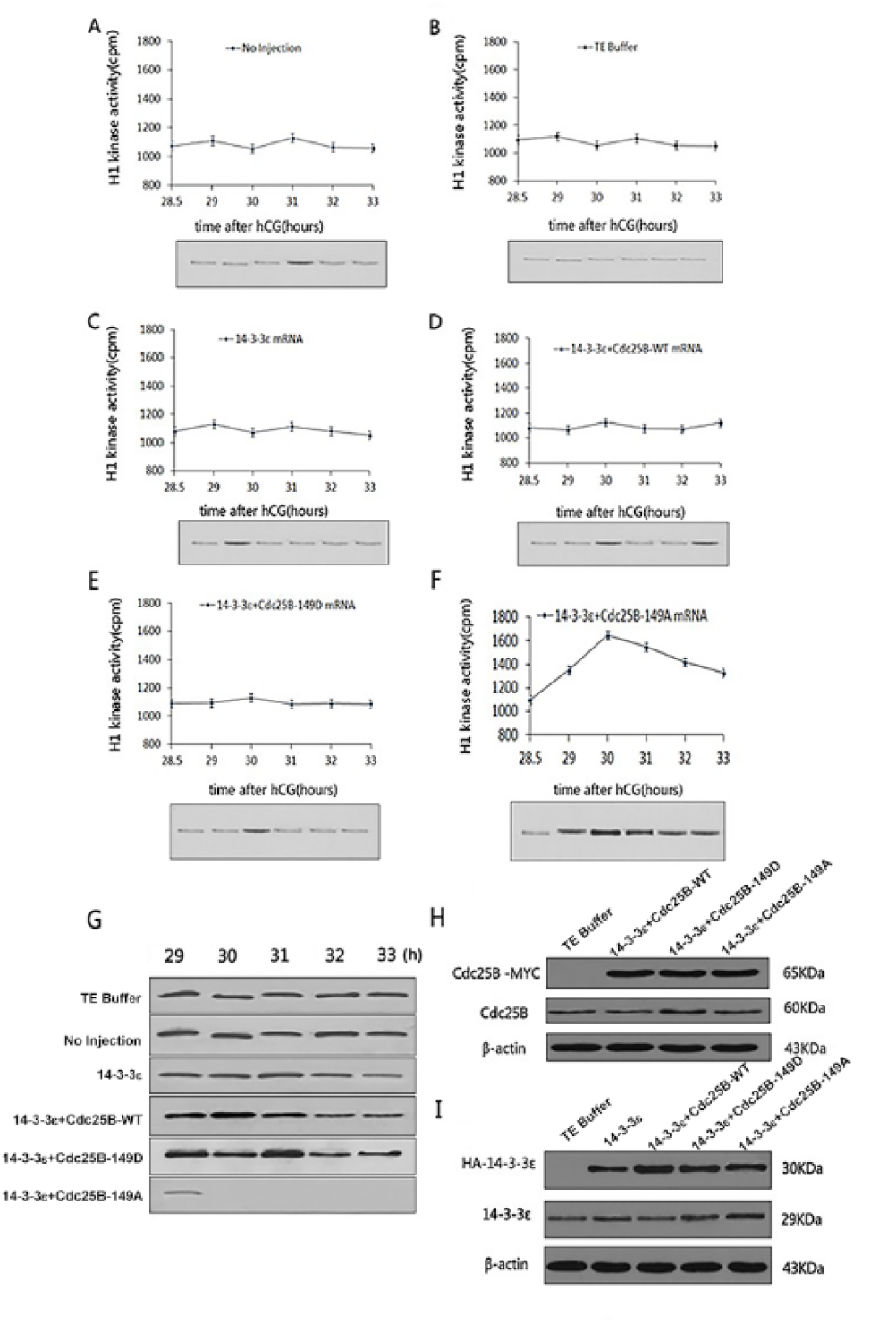
Variation of MPF activity and Cdc2-Tyr15 phosphorylation status in the co-injected Cdc25B-Ser149A and 14-3-3ε mRNA group and other mRNA-injected groups. Effects of microinjection with 14-3-3ε mRNA solely or co-injection with Cdc25B-WT mRNA, Cdc25B-Ser149A mRNA or Cdc25B-Ser149D mRNA and 14-3-3ε mRNA on mitotic resumption of mouse fertilized eggs. Experiments were performed in the continued presence of 2mmol/l dbcAMP (A–G). (A) MPF activity of no injection group. (B) MPF activity of TE buffer group. (C) MPF activity of co-injection with 14-3-3ε mRNA. (D) MPF activity of co-injection with 14-3-3ε mRNA and Cdc25B-WT mRNA. (E) MPF activity of co-injection with 14-3-3ε mRNA and Cdc25B-Ser149D mRNA. (F) MPF activity of co-injection with 14-3-3ε mRNA and Cdc25B-Ser149A mRNA. For each point, five eggs were collected, and MPF activity was examined by scintillation counting and autoradiography. One-way analysis of variance followed by a Least Significant Difference (LSD) test was used to evaluate the differences in the MPF activity assay. Bars represent means ± S.D of three independent experiments. (G) Western blot analysis of the phosphorylation status of Cdc2-Tyr15 in the various mRNA injection groups. The eggs were collected at 29, 30, 31,32 and 33 h after the hCG injection. A total of 200 eggs were loaded onto each lane in Western blot. (H) Western analysis of Cdc25B-MYC and endogenous Cdc25B expression. The mouse fertilized eggs co-injected with Cdc25B-WT mRNA, Cdc25B-Ser149D mRNA or Cdc25B-Ser149A mRNA and 14-3-3ε mRNA were collected 4 h after injection and their proteins were immunoblotted with anti-myc, anti-Cdc25B or anti-β-actin antibody. Bars represent means ± S.D of three independent experiments. (I) Western analysis of HA-14-3-3ε and endogenous 14-3-3ε expression. The mouse fertilized eggs co-injected with Cdc25B-WT mRNA, Cdc25B-Ser149D mRNA or Cdc25B-Ser149A mRNA and 14-3-3ε mRNA were collected 4 h after injection and their proteins were immunoblotted with anti-HA, anti-14-3-3ε or anti-β-actin antibody. Bars represent means ± S.D of three independent experiments.

**Figure 4.**
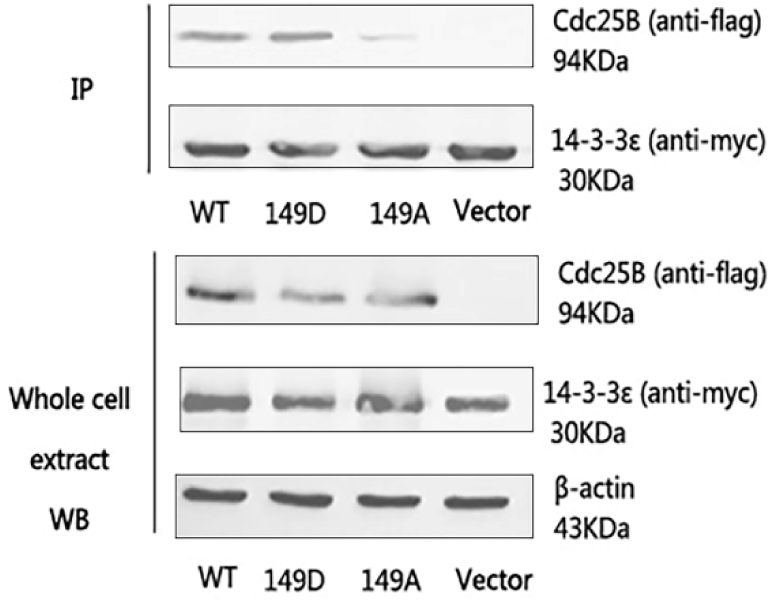
14-3-3ε was immunoprecipitated with anti-HA beads followed by Western blotting to detect Cdc25B binding with 14-3-3ε. Western blot analysis demonstrating the expression of Myc-14-3-3ε and Flag-Cdc25B in the cell lysates (Input) used to immunoprecipitate Cdc25B protein shown in (IP), an anti-β-actin antibody was used to determine equal loading of the gel.

### Co-localization of endogenous expressed 14-3-3ε and Cdc25B in fertilized eggs

Using indirect immunofluorescence, we observed the co-localization of endogenous Cdc25B and 14-3-3ε at each phase of cell cycle in fertilized mouse eggs. We checked 30 different eggs from G1, S, early G2, late G2 and M phases, respectively, and all showed the same pattern of immunofluorescent staining. As shown in Figure 5, green fluorescent Cdc25B signals and red fluorescent 14-3-3ε signals were co-localized primarily in the cytoplasm at G1 and S phases, respectively (Figure 5A and B). Green fluorescent Cdc25B signals and red fluorescent 14-3-3ε signals were observed in the cytoplasm, the partial green fluorescent Cdc25B signals translocated to the nucleus of mouse embryos at early G2 phase (Figure 5C). However, the Cdc25B signals in the nucleus was significantly enhanced at late G2 phase (Figure 5D), whereas 14-3-3ε signals remained in the cytoplasm at late G2 phase (Figure 5C and 5D). At M phase The green fluorescent Cdc25B signals and red fluorescent 14-3-3ε were evenly distributed in the cytoplasm of the whole cell again (Figure 5E). These results indicated that the shuttling of Cdc25B is accompanied with G2/M transition.

**Figure 5.**
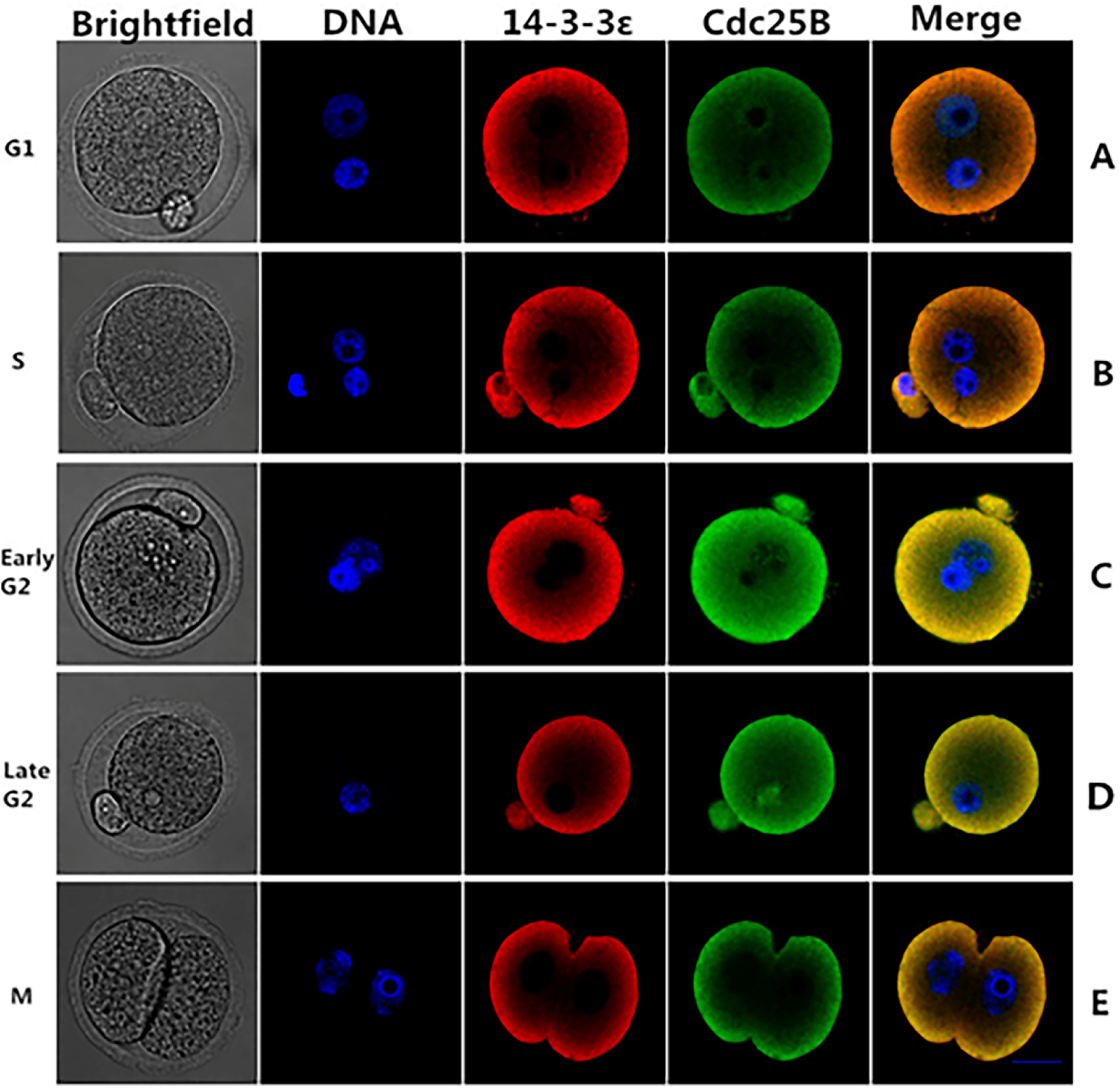
Co-localization of endogenous 14-3-3ε and Cdc25B in fertilized eggs. The representative cells were fixed, permeabilized and immunolabeled for confocal microscopy at different phases. (A, B) The green fluorescent Cdc25B signals and red fluorescent 14-3-3ε signals were co-localized primarily in the cytoplasm at G1 and S, respectively. (C) the partial green fluorescent signals Cdc25B were translocated to the nucleus in early G2 phase and red fluorescent 14-3-3ε signals were detected in the cytoplasm. (D) red fluorescent 14-3-3ε signals remained in the cytoplasm, but the nuclear accumulation of green fluorescent Cdc25B signals were observed at late G2 phase. (E) The green fluorescent Cdc25B signals and red fluorescent 14-3-3ε signals were evenly distributed in the cytoplasm again at M phase. Scale bar=20μm.

### Co-localization of exogenously expressed 14-3-3ε and Cdc25B in fertilized eggs

To confirm the effect of the Ser149 phosphorylation on subcellular localization of Cdc25B and 14-3-3ε, pEGFP-Cdc25B-WT or pEGFP-Cdc25B-Ser149A was co-injected with pRFP-HA-14-3-3ε into fertilized mouse eggs of G1 phase (19h after hCG injection), and then the microinjected eggs were transferred into M16 medium containing 2 mmol/L dbcAMP to allow protein expression. As shown in Figure 6, when the two groups of fertilized mouse eggs entered early G2 phase, green fluorescent Cdc25B (WT/Ser149A) signals and red fluorescent 14-3-3ε signals were co-localized in the cytoplasm of mouse embryos at early G2 phase (Figure 6A and B). In contrast, the green fluorescent signals of Cdc25B-Ser149A were translocated to the nucleus, whereas Cdc25B-WT signals were still retained in the cytoplasm of mouse fertilized egg without nucleus accumulation at late G2 phase (Figure 6C and D).14-3-3ε signals were detected primarily in the cytoplasm in both groups at late G2 phase (Figure 6C and D). Moreover, localization was independent of the amount of recombinant protein expressed and of the tag used to detect the expressed proteins [28]. These results suggest that Cdc25B cannot transfer to the nucleus when Cdc25B-Ser149 is phosphorylated and Cdc25B-Ser149 was indeed critical for the subcellular localization of Cdc25B.

**Fiigure 6.**
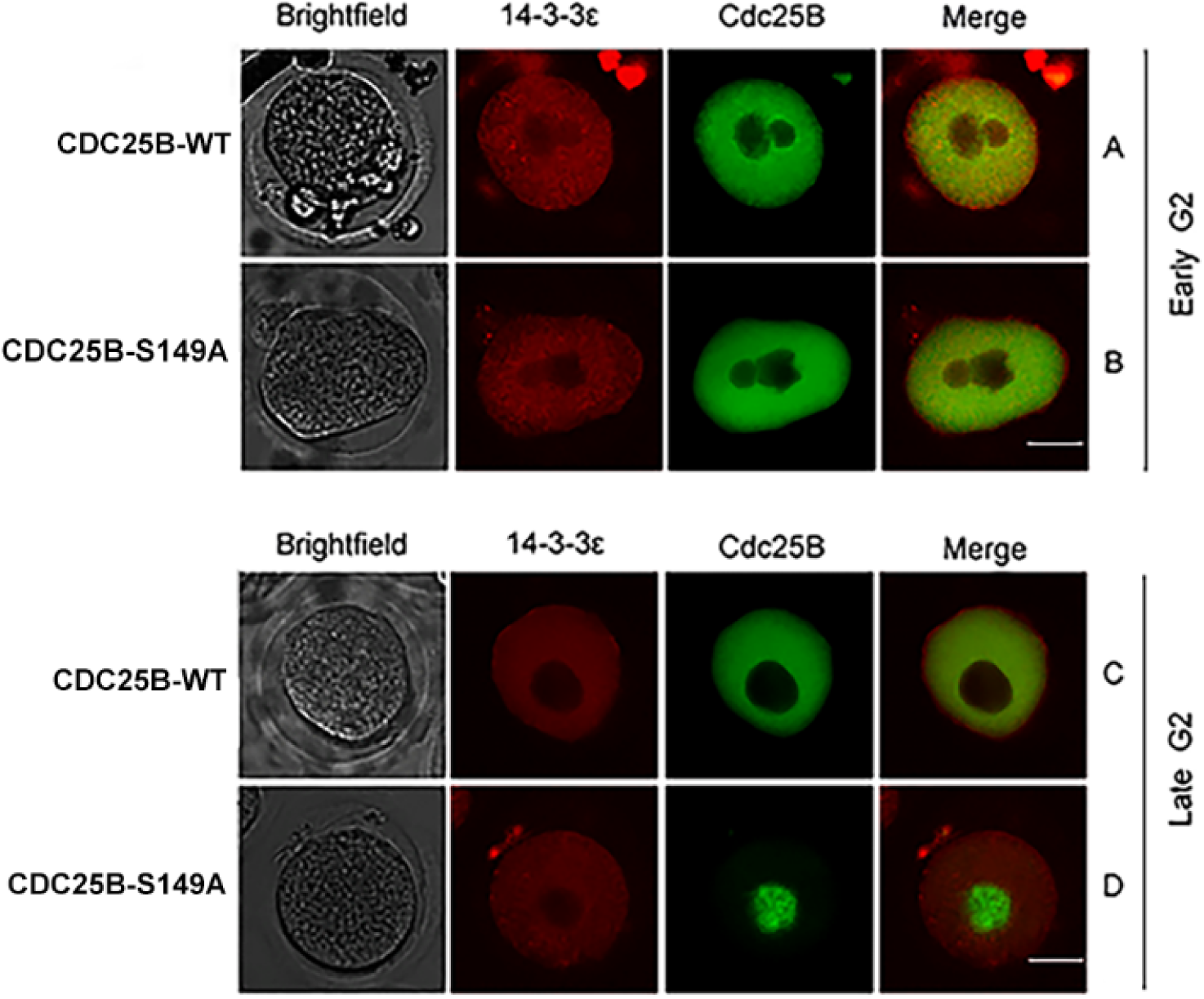
Co-localization of exogenously expressed 14-3-3ε and Cdc25B in fertilized eggs. (A, B) green fluorescent Cdc25B(WT/Ser149A) signals and red fluorescent 14-3-3ε signals were co-localized primarily in the cytoplasm at early G2 phase. (C) green fluorescent Cdc25B-WT signals were detected in the cytoplasm at late G2 phase. (D) Cdc25B-Ser149A translocated to the nucleus whereas 14-3-3ε localized predominantly in the cytoplasm at late G2 phase. Scale bar =20μm.

## Discussion

PKA, also known as 3’, 5’-cyclic phosphate-dependent protein kinase A, is the most common and earliest discovered member of the protein kinase superfamily. It regulates cell growth, reproduction, differentiation and specific protein expression by phosphorylating specific protein serine/threonine residues [30–34]. It has been demonstrated that in mammals, Cdc25B acts as a direct downstream substrate for PKA and regulates the progression of the mouse one-cell embryos by phosphorylation or dephosphorylation of Ser149 and Ser321 of Cdc25B [9, 24, 25]. Our previous LC-MS/MS analysis showed that PKA can phosphorylate the Ser149, Ser229, and Ser321 sites of Cdc25B, corresponding to the phosphorylation sites predicted by the Scansite software[8, 28]. Our previous studies have shown that prophase I arrest in mouse oocytes depends on phosphorylation of Cdc25B-Ser321 (not Cdc25B-Ser149) mediated by PKA and Cdc25B binding to 14-3-3ε protein contributes to the maintenance of prophase I arrest in the oocytes[28]. Cui et al. [7] showed that 14-3-3ε binding to Cdc25B-Ser321 phosphorylated by PKA induces mitotic arrest at one-cell stage by inactivating MPF in mouse fertilized eggs. Meanwhile, we have confirmed that Ser149 of Cdc25B was phosphorylated in the G1 and S stage and dephosphorylated in the G2 and M stage [8]. Above data strongly suggest that at high cAMP levels, activated PKA phosphorylates Cdc25B on Ser149 and Ser321, the inhibitory phosphorylation of Cdc25B induces mitotic arrest at one-cell stage in mouse fertilized eggs or maintenance of prophase I arrest in the oocytes by negative regulating MPF.

14-3-3 proteins are a highly conserved, ubiquitously expressed family of proteins, including seven known isoforms in mammalian species (β, γ, ε, η, δ, τ, ζ). These proteins associate with many intracellular proteins involved in a variety of cellular processes including regulation of the cell cycle, metabolism and protein trafficking [35–37]. The 14-3-3 protein provides a functional basis site for binding to target proteins by forming a homodimer or heterodimer, consequently, acting as molecular chaperone or intermediate [38].

In this study, we explored whether one-cell stage mouse fertilized eggs express different 14-3-3 isoforms. The data showed that one-cell stage fertilized mouse eggs express all seven 14-3-3 isoforms and the relative expression of 14-3-3ε and 14-3-3ζ mRNAs were higher than that of the other five isoforms. The protein expression of 14-3-3ε gradually increased from G1 to G2 and decreased slightly in M phase, consistent with its mRNA expression. Consistent with our results, Santanu De and his colleagues [27] have reported that mouse mature metaphase II-arrest eggs express all seven 14-3-3 isoforms. Contrary to our results, Cui C and her colleagues [9] have reported that only 14-3-3ε existed in G1 phase of fertilized mouse eggs. Here, we will focus our discussion on the effect of 14-3-3ε on G2/M transition of one-cell stage mouse fertilized eggs. In this study, we provide experimental evidence for the interaction and co-localization of 14-3-3ε and Cdc25B-Ser149 in G2/M transition of mouse fertilized eggs. Our indirect immunofluorescence experiments revealed a restricted cytoplasmic co-localization of 14-3-3ε and Cdc25B at G1, S, early G2 phases whereas nuclear localization of Cdc25B with cytoplasmic localization of major 14-3-3ε at late G2 phase, in agreement with previous reports [39, 40].

Our previous study demonstrated that overexpression of Cdc25B-Ser321A alone in one-cell stage mouse fertilized eggs can induce Cdc2-Tyr15 dephosphorylation and mitosis resumption much more efficiently than overexpression of Cdc25B-WT or Cdc25B-Ser321D [7]. These results suggest that 14-3-3ε may inhibit Cdc25B activity by binding to phosphorylated Ser321, eventually fails to activate MPF for further mitosis resumption. Here our findings demonstrated that co-expression of 14-3-3ε and Cdc25B-Ser149A in one-cell stage mouse fertilized eggs was able to initiate mitosis, whereas co-expression of Cdc25B-WT or Cdc25B-Ser149D and 14-3-3ε was unable to initiate mitosis. In addition, overexpression of 14-3-3ε alone cannot affect the G2/M transition of mouse fertilized eggs. Thus, the ability of the co-expression of 14-3-3ε to inhibit Cdc25B-dependent activation of MPF is likely related to 14-3-3ε binding of Cdc25B-WT, which is defective in the Ser149A mutant. Studies have shown that the Cdc25B-Ser149 is phosphorylated at the G1 and S phases in the fertilized mouse eggs, whereas no phosphorylation of Cdc25B-Ser149 was observed at the G2 and M phases in vivo, suggesting that unphosphorylated Cdc25B-Ser149A is required for activating MPF [9, 40]. These findings further consolidated our conclusion that 14-3-3ε binds to phosphorylated Ser149 and inhibits Cdc25B activity. It is reported that the Cdc25B-Ser149A functions similarly to Cdc25B-Ser321A but is much less efficient than the Cdc25B-Ser149A/Ser321A mutant[8], suggesting that Ser149 and Ser321of Cdc25B may be the direct targets of PKA in mammals and synergistically regulate the early development of fertilized mouse eggs. About the specific interaction mechanism of 14-3-3ε and Cdc25B, these data strongly suggest that Ser149 is phosphorylated by PKA that may provide a docking site for consequent 14-3-3ε binding, which in turn covers the NES of Cdc25B, thereby causing nuclear exclusion of the protein without affecting its phosphatase activity. Our finding of the co-localization of exogenously 14-3-3ε and Cdc25B and the immunoprecipitation further demonstrates that when Ser149 of Cdc25B is mutated to an unphosphorylated residue Ala, the 14-3-3ε binding is canceled and signal of Cdc25B nucleus export is disrupted. Therefore, we speculate the localization of Cdc25B is likely regulated by the binding with 14-3-3ε. To sum up, the Ser149 and Ser321 residues of Cdc25B may be potential targets for 14-3-3ε to participate in mitosis of fertilized mouse eggs in mammalian cells and play an important role in G2/M transition of fertilized mouse eggs and in the subcellular localization of Cdc25B.

## Author Contributions

**He Wenning:** conceptualization; formal analysis ; investigation; writing— review & editing; supervision. **Pang Haiyao:** conceptualization; methodology; formal analysis; investigation; writing— review & editing; visualization. **Liu Ru:** methodology; resources. **Zhang Huiling:** validation; formal analysis; data curation. **Guo Wenxiu:** validation; formal analysis; data curation. **Feng Shaoqing:** investigation. **Hou Yanjun:** writing—original draft. **Meng Jun:** methodology; conceptualization; original draft; funding acquisition; project administration.

## Acknowledgments

This work was supported by the National Natural Science Foundation of China (81360109 and 81660267).

